# Discovery and characterization of variance QTLs in human induced pluripotent stem cells

**DOI:** 10.1101/424192

**Authors:** Abhishek K. Sarkar, Po-Yuan Tung, John D. Blischak, Jonathan E. Burnett, Yang I. Li, Matthew Stephens, Yoav Gilad

**Affiliations:** Department of Human Genetics, University of Chicago, Chicago, Illinois, USA; Department of Medicine, University of Chicago, Chicago, Illinois, USA; Department of Statistics, University of Chicago, Chicago, Illinois, USA

## Abstract

Quantification of gene expression levels at the single cell level has revealed that gene expression can vary substantially even across a population of homogeneous cells. However, it is currently unclear what genomic features control variation in gene expression levels, and whether common genetic variants may impact gene expression variation. Here, we take a genome-wide approach to identify expression variance quantitative trait loci (vQTLs). To this end, we generated single cell RNA-seq (scRNA-seq) data from induced pluripotent stem cells (iPSCs) derived from 53 Yoruba individuals. We collected data for a median of 95 cells per individual and a total of 5,447 single cells, and identified 241 mean expression QTLs (eQTLs) at 10% FDR, of which 82% replicate in bulk RNA-seq data from the same individuals. We further identified 14 vQTLs at 10% FDR, but demonstrate that these can also be explained as effects on mean expression. Our study suggests that dispersion QTLs (dQTLs) which could alter the variance of expression independently of the mean can have larger fold changes, but explain less phenotypic variance than eQTLs. We estimate 424 individuals as a lower bound to achieve 80% power to detect the strongest dQTLs in iPSCs. These results will guide the design of future studies on understanding the genetic control of gene expression variance.

**Author summary:** Common genetic variation can alter the level of average gene expression in human tissues, and through changes in gene expression have downstream consequences on cell function, human development, and human disease. However, human tissues are composed of many cells, each with its own level of gene expression. With advances in single cell sequencing technologies, we can now go beyond simply measuring the average level of gene expression in a tissue sample and directly measure cell-to-cell variance in gene expression. We hypothesized that genetic variation could also alter gene expression variance, potentially revealing new insights into human development and disease. To test this hypothesis, we used single cell RNA sequencing to directly measure gene expression variance in multiple individuals, and then associated the gene expression variance with genetic variation in those same individuals. Our results suggest that effects on gene expression variance are smaller than effects on mean expression, relative to how much the phenotypes vary between individuals, and will require much larger studies than previously thought to detect.

## Introduction

Robustness, or the ability to maintain a stable phenotype despite genetic mutations and environmental perturbations, is an important property of many key biological processes, such as those underlying embryogenesis and development [1, 2]. Conversely, evolvability, or the ability to generate heritable phenotypic variation, is a fundamental requirement of evolutionary processes [3]. A long-standing question in genetics, therefore, is how the balance between these two seemingly opposite processes has been fine-tuned [4].

We still have a relatively poor understanding of how robustness is achieved, especially at the molecular level. In model organisms, robustness and evolvability can be studied using experimental evolution approaches. These approaches typically quantify robustness as the change in trait variation after applying an experimental perturbation [5, 6]. However, in such experiments the phenotypic outcomes, rather than the underlying mechanisms of robustness, are measured. Moreover, experimental evolution studies have almost always considered population-average measurements of phenotypes using entire organisms, tissues, or cell cultures, with few exceptions [7, 8]. To truly understand how robustness and evolvability are established and encoded in the genome, we need to consider phenotypic variation across individual cells [9], and connect it to genetic variation, an approach termed “noise genetics” [10].

Using the yeast *Saccharomyces cerevisiae* as a model system, studies have shown that heterogeneity in the expression of certain genes across cells is highly heritable and placed under complex genetic control, suggesting that the level of noise in gene regulation may also differ between individuals of multicellular organisms depending on their genetic background [11]. Follow-up studies further demonstrated that gene expression noise mediated by promoter variants can provide a fitness benefit at times of environmental stress in yeast, highlighting the direct role of genetically controlled stochastic cell-cell variation in evolutionary robustness [12]. However, the genetic and molecular circuits that lead to robustness remain largely uncharacterized in mammals.

Here, we take an unbiased, genome-wide approach to identify quantitative trait loci associated with gene expression variance across cells (vQTLs). We study human induced pluripotent stem cells (iPSCs), which offer a homogeneous population of cells allowing a relatively simple statistical model. Investigating iPSCs also provides the possibility to study gene expression variance across cells during differentiation in follow-up studies. To directly measure the mean and variance of gene expression within cell populations as phenotypes, we generated single cell RNA-seq (scRNA-seq) data from cells derived from multiple individuals.

## Results

### Sample collection and quality control

Using the Fluidigm C1 platform, we isolated and collected scRNA-seq from 7,585 single cells from iPSC lines of 54 Yoruba in Ibadan, Nigeria (YRI) individuals. We used unique molecular identifiers (UMIs) to tag RNA molecules and account for amplification bias in the single cell data [13]. To estimate technical confounding effects without requiring separate technical replicates, we used a mixed-individual plate study design (Fig 1A). The key idea of this approach is that having observations from the same individual under different confounding effects and observations from different individuals under the same confounding effect allows us to distinguish the two sources of variation [14].

**Figure 1:**
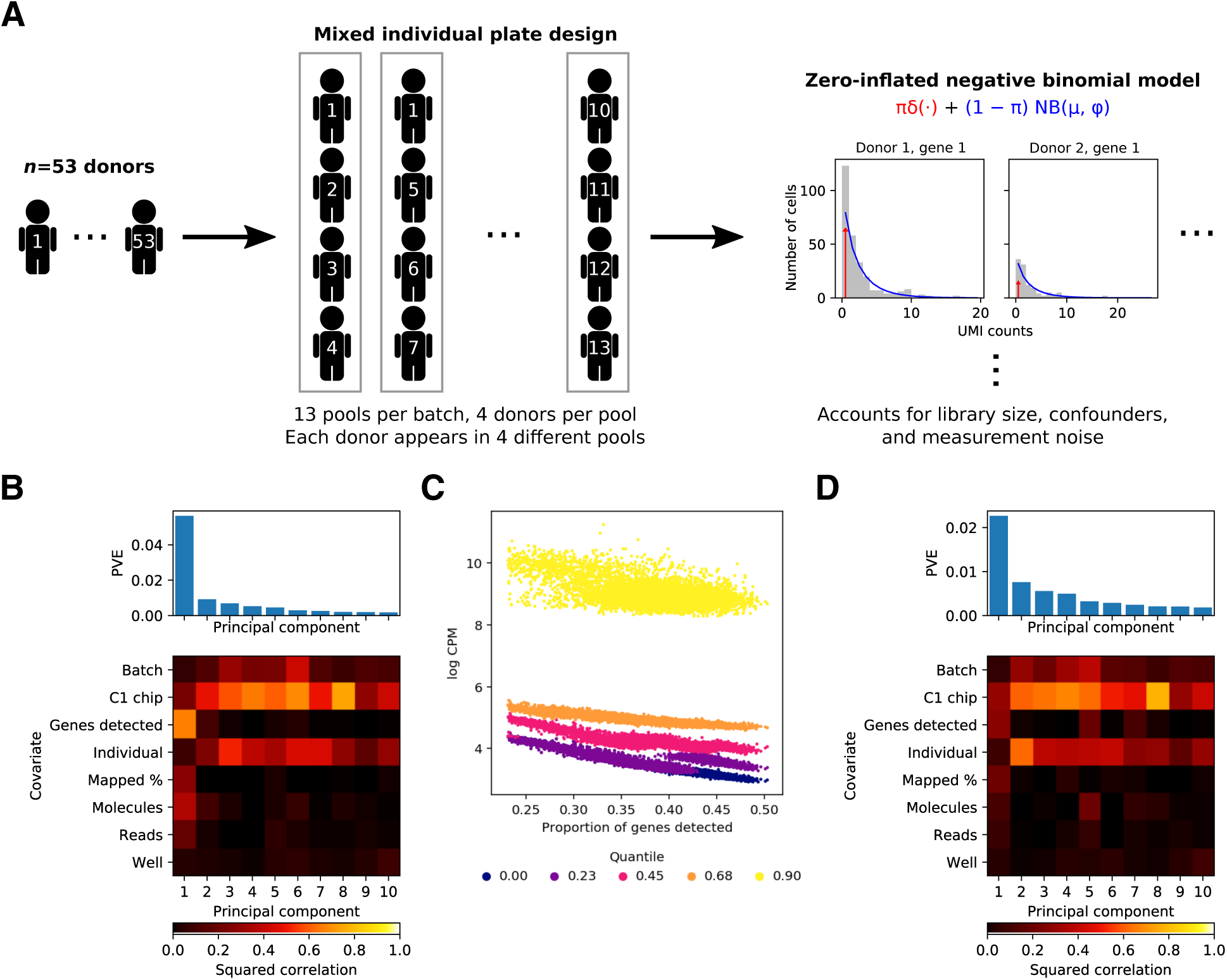
Study design and quality control. (A) We used a mixed-individual plate design to be able to distinguish technical effects from biological effects of interest, and used a zero-inflated negative binomial model to fit the distribution of the data, accounting for technical confounders. (B) Proportion of variance explained (PVE; top) and a heatmap of the correlations between the top 10 principal components of gene expression and observed technical covariates (bottom). (C) Dependence of the distribution of gene expression against gene detection rate (proportion of genes with at least one molecule detected) for each sample. Each vertical slice is a single cell (according to the gene detection rate). For each cell, there are 5 points, corresponding to the (0,0.25,0.5, 0.75,1) quantiles of non-zero log CPM values observed for that cell. (D) PVE and correlation between principal components and observed covariates after correcting for gene detection rate.

We excluded data from one individual (NA18498) with evidence of contamination, then filtered poor quality samples as previously described [14]. After quality control, we analyzed the expression of 9,957 protein-coding genes in a median of 95 cells per individual in 53 individuals (total of 5,597 cells; Fig S1).

To ensure that our measurements are comparable across samples, we first sought to assess the impact of observed technical variation on the data and to identify unobserved technical confounders. To this end, we performed principal components analysis (PCA) on the matrix of log counts per million (log CPM).

We found that across samples, the loading on the top principal component (PC) was correlated with gene detection rate (the proportion of genes with at least one molecule detected), but not with the biological variable of interest (individual) or the expected technical confounders (batch and C1 chip; Fig 1B). Indeed, as previously reported [15], the entire distribution of observed log CPM (over all genes) varies across samples, and appears to be associated with the gene detection rate (Fig 1C). After accounting for gene detection rate (Methods), the top PCs were correlated with individual, batch, and C1 chip, as expected (Fig 1D).

### Estimating gene expression mean and variance

We developed a method to estimate the mean and variance of gene expression across cells for each gene in each individual (Fig 1A; Methods). Briefly, for each individual and each gene, our method uses maximum likelihood to fit a zero-inflated negative binomial distribution (ZINB) to the observed UMI counts across cells, and derives the mean and variance of gene expression from the estimated model parameters. When fitting the ZINB model the method controls for technical confounders (e.g. C1 chip) and library size, and when deriving the mean and variance it accounts for Poisson measurement noise in the UMI counts [16, 17]. These desirable features would not be achieved by directly computing the sample mean and variance of either the UMI counts or log CPM.

To evaluate the accuracy of the method, we simulated data from the model and compared the estimated parameters, as well as the derived mean and variance, to the true values used to generate the data. We fixed the number of cells and number of molecules detected per cell to the median of those values in our observed data, and varied the ZINB parameters. Assuming that mean expression is high enough, we found the method produces accurate estimates of the underlying negative binomial parameters, but not the zero inflation parameter (Fig S2). Despite not accurately estimating the zero inflation parameter, the method still produces accurate estimates of the derived mean and variance for genes that are expressed at intermediate to high levels.

We thus applied our method to the observed data, correcting for batch and C1 chip. Importantly, we did not correct for gene detection rate, reasoning that the dependence on gene detection rate is only an artifact introduced by analyzing log CPM. To ensure our method would provide accurate estimates for the observed data, we filtered genes based on their median expression level across samples. We then asked what proportion of variance in the expression data was explained by technical confounders. To do so, we estimated the reduction in residual variance when including the confounders in the model, and found technical confounding explains 16% of the observed variance on average across all genes and individuals. Our results emphasize that careful experimental design as well as careful statistical modeling are required to robustly map effects on gene expression variance across cells.

### Quantitative trait locus mapping

Previous studies have shown a clear relationship between the mean and variance of gene expression [18, 19]; therefore, apparent genetic effects on the variance could potentially be explained by effects on the mean. In our model, the mean-variance relationship is controlled by a single dispersion parameter per gene per individual. We sought to directly map QTLs which could alter the variance independently of altering the mean by using the estimated dispersion parameter as a quantitative phenotype. However, we found zero dispersion QTLs (dQTLs) using this approach.

Alternative approaches to decouple the mean-variance relationship include using the coefficient of variance (CV; ratio of standard deviation to mean) or Fano factor (ratio of variance to mean) as quantitative phenotypes. However, prior work shows these quantities have predictable relationships with the mean, and therefore effects could still be explained away [14, 19]. Therefore, we proceeded to map eQTLs, variance QTLs (vQTLs), CV-QTLs, and Fano-QTLs, and then asked whether we could discover variance effects which could not be explained as effects on mean expression.

We found 241 eQTLs, 14 vQTLs, 2 CV-QTLs, and 0 Fano-QTLs (FDR 10%). To validate the eQTLs, we estimated the replication rate against eQTLs discovered in bulk RNA-seq from the same iPSC lines [20]. We found that 82% of the single cell eQTLs replicate in the matched bulk data (Fig 2A), and 80% of bulk eQTLs replicate in the single cell data. However, we found 1,390 eQTLs (FDR 10%) using all of the individuals in the bulk RNA-seq study (*n* = 58), and still recovered 1,136 eQTLs (FDR 10%) after subsampling to *n* = 53. Our results therefore suggest that eQTL discovery in scRNA-seq (as opposed to replication of previously discovered eQTLs) loses power compared to equal-sized studies in bulk RNA-seq, likely due to increased experimental noise.

**Figure 2:**
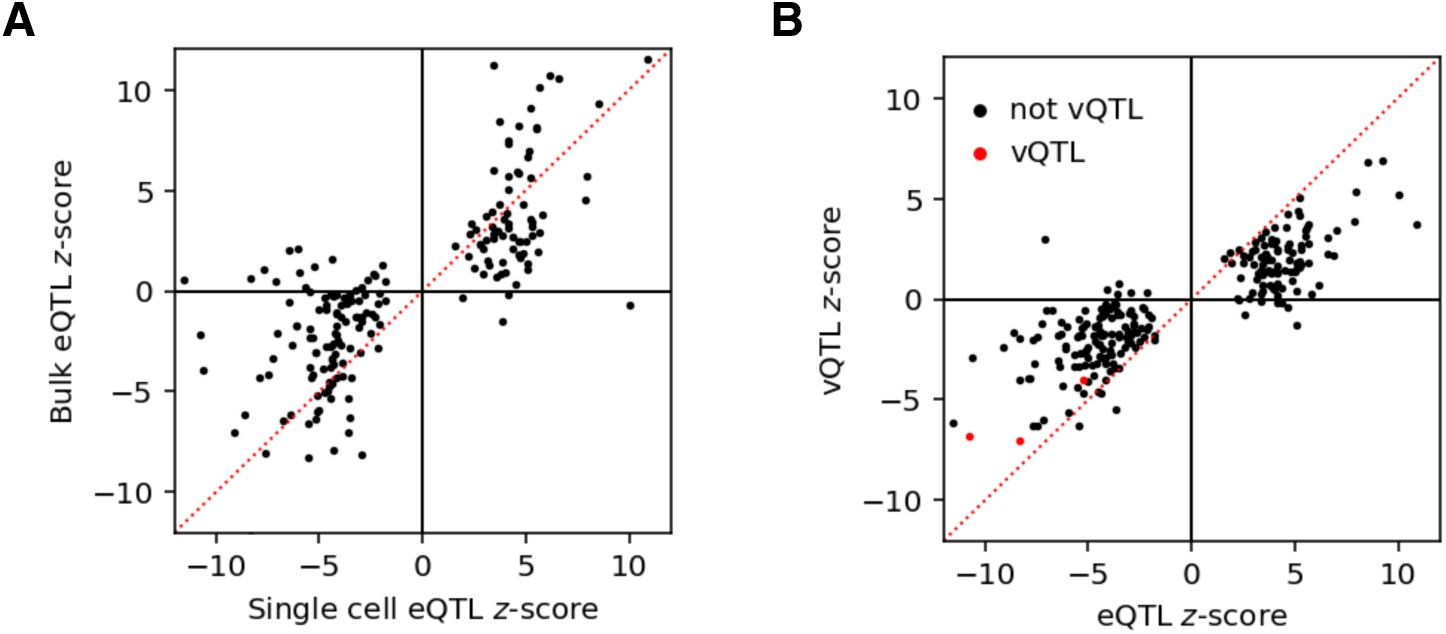
Discovery and overlap of expression QTLs and variance QTLs. (A) *z*-scores for expression QTL (eQTL) SNP-gene pairs discovered in pooled single cell RNA-seq data against *z*-scores of the same SNP-gene pairs in matched bulk RNA-seq data. (B) In the single cell data, *z*-scores for 241 eQTL SNP-gene pairs (FDR 10%) against variance QTL (vQTL) *z*-scores of the same SNP-gene pairs. vQTL *z*-scores are stratified based on whether the gene was discovered as a vQTL at FDR 10%.

We then sought to directly explain away vQTLs as eQTLs by regressing out the mean from the variance. Treating the residuals from the regression as the phenotype, we recovered zero vQTLs. Similarly, after regressing out the mean from the coefficient of variation, we recovered zero CV-QTLs. These results suggest the significant variance effects detected in this study are all likely to be explained as effects on mean expression.

### Power analysis

Our goal in this study was to find QTLs which alter the variance of gene expression independently of altering the mean expression. Under our model, these QTLs should explain variation in the dispersion parameter across individuals; however, we failed to find dQTLs. Further, all of the vQTLs we were able to identify could be explained by mean effects. In contrast, we were able to discover eQTLs, but fewer than expected based on bulk RNA-seq in matched samples.

To understand why we failed to discover dQTLs, and why we discovered fewer eQTLs than expected, we first derived the power function in terms of effect size (log fold change), sample size, noise ratio (ratio of measurement error variance to phenotypic residual variance), and significance level (Methods). We then sought to estimate the distribution of QTL effect sizes and the typical noise ratio, for the both mean expression and dispersion.

To estimate the distribution of QTL effect sizes, we fit a flexible unimodal distribution for the true effect sizes which maximizes the likelihood of the observed effect sizes and standard errors [21]. Surprisingly, we found that dQTL effects could be larger than eQTL effects (Fig S3). For example, we estimate that the 99^th^ percentile eQTL effect size is 0.023, but is 0.085 for dQTLs. Given this result and the power function we derived, there are two possible explanations for why we still failed to find dQTLs: (1) the noise ratio of dispersion is large (measurement error reduced power), or (2) the residual variance of dispersion is large (genetic variation explains little phenotypic variance).

To estimate the typical noise ratio, we developed a two-step procedure to estimate the measurement error variance and residual variance per gene (Methods). Briefly, in our approach we have one measurement error variance per individual, per gene, which equals the sampling variance of our ZINB model. To estimate each error variance, we used non-parametric bootstrapping. To estimate the measurement error variance for each gene, we took the median of the estimated measurement error variances across individuals. To estimate the residual variance for each gene, we fit a flexible unimodal distribution for the true phenotypes which maximizes the likelihood of the observed phenotypes and measurement errors, and estimated the variance of the posterior mean true phenotypes.

Using our approach, we estimated that the typical noise ratio of the dispersion is 3.0, compared to 8.0 for the mean (Figure S4). This result suggests that we did not fail to find dQTLs only due to measurement error, because the noise ratio was lower for dQTLs than for eQTLs. As a reference point, a noise ratio equal to 1 has the same impact on power to detect a QTL as cutting the sample size in half, explaining why out study lost power to detect eQTLs. We found that the typical phenotypic standard deviation of dispersion is 7.2 fold larger than that of the mean expression, suggesting we failed to find dQTLs because the effect sizes of dQTLs (relative to phenotypic standard deviation) are smaller than the effect sizes of eQTLs.

We finally asked how much power our current study had to detect the 99^th^ percentile dQTL effect size, assuming the typical noise ratio estimated above. We found that our study had only 0.02% power to detect that effect size at Bonferroni–corrected level *α* = 5 × 10^−6^ (Figure 3). Fixing the typical noise ratio, we estimate 1,702 individuals would be required to achieve 80% power. As a lower bound, we estimate 424 individuals would be required assuming no measurement error. Overall, our results suggest a much larger study, both in terms of number of individuals and number of cells per individual, would be required to detect the strongest dQTLs in iPSCs.

**Figure 3:**
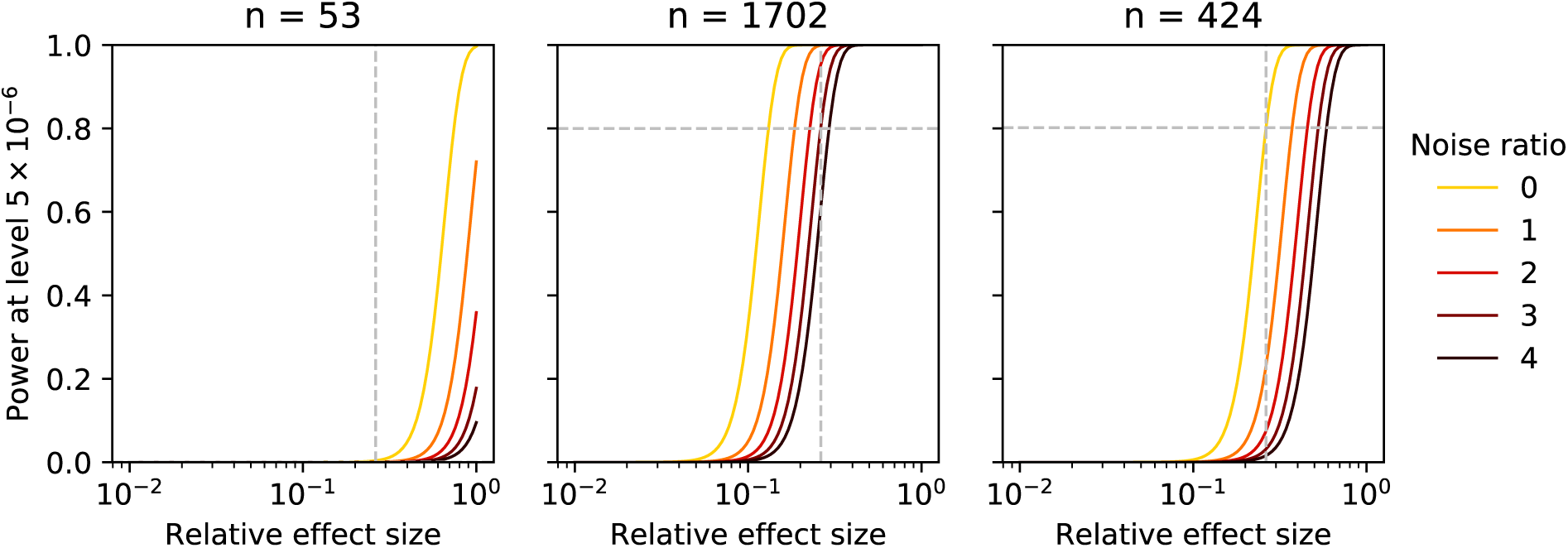
Power to detect dispersion QTLs. Power is a function of effect size (relative to phenotypic standard deviation), sample size, noise ratio, and significance level. Gray lines indicate the 99^th^ percentile of dispersion effect sizes relative to the typical phenotypic standard deviation, and the power achieved to detect an effect of that size at the typical noise ratio. Power curves are computed for the current sample size (left), the sample size required to achieve 80% power for that effect size fixing the number of cells per individual (center), and the minimum sample size assuming no measurement error (right).

## Discussion

Individual cells must tolerate both external and internal perturbations arising from the environment or mutations. It has long been argued that this outcome of robustness is an inherent property of biological systems [22], and arises from natural selection [23, 24]. Robustness is especially critical in the context of cell fate transitions during differentiation [25]. Other dynamic physiological processes must also be robust, and as a result, loss of robustness is associated with clinically relevant phenotypes and complex genetic disease [26, 27].

Cells maintain their identity and other phenotypes despite perturbations because of the robust regulation of key sets of genes [28]. We hypothesized that QTLs could disrupt the mechanisms underlying robust regulation, and therefore reveal new insights into the genetic regulation of differentiation and disease.

To investigate this hypothesis, we directly observed gene expression variance across multiple individuals using scRNA-seq, and sought to identify QTLs which could alter the variance of gene expression across cells within a single individual, independently of altering the mean expression. However, we failed to discover such QTLs, and demonstrated that QTLs which are associated with the variance of gene expression can be explained by effects on mean expression. We found that relative to the phenotypic standard deviation, effects on the dispersion are smaller than effects on the mean, partially explaining why this study failed to find them.

Our results do not rule out genetic effects on variance independent of mean effects, due to two main limitations of our analysis. First, our estimated distributions of effect sizes are based on an empirical Bayes estimate of the underlying effect sizes, given the observed effect sizes. Our results in simulation and observed data suggest the observed effect sizes may be not be accurately estimated given the size of the current study. Therefore, the empirical Bayes estimate may not accurately reflect the true distribution of effect sizes. However, we chose to bias the estimation procedure towards putting prior mass on zero, so our estimates of effect sizes are conservative. Additionally, our estimates may not generalize beyond iPSCs, because the distribution of dispersion effect sizes could vary across cell types and conditions.

Second, we took a modular approach to map QTLs in this study: (1) we estimated parameters for each individual using only the scRNA-seq data, and then (2) we mapped QTLs using phenotypes derived from the estimated parameters. An alternative approach would be to include genotype in the count model for the data, and jointly learn the mean, dispersion, proportion of excess zeros, and genetic effect sizes for mean and dispersion. Such an approach could borrow information across cells with common genotypes to improve power, holding the experiment size fixed. However, further development will be needed to efficiently fit the models at QTL mapping scale.

We stress that our power calculation is only a rough guideline for designing QTL mapping studies using scRNA-Seq. We based our calculations on typical values of the noise ratio for the mean expression and dispersion, and chose a conservative significance level. However, we found considerable variation in the noise ratio across genes, suggesting that our results may not generalize even across genes. Overall, our results suggest that the technical noise introduced by scRNA-seq greatly reduces the power to discover eQTLs. Our results also suggest that, for iPSC lines, dramatically larger studies will be required to map both eQTLs and dQTLs from scRNA-seq.

## Materials and methods

### Sample collection and quality control

We cultured YRI iPSCs [20] in feeder-free conditions for at least ten passages in E8 medium (Life Technologies) [29]. We collected cells using the C1 Single-Cell Auto Prep IFC microfluidic chip (Fluidigm). We used a balanced block-incomplete design to randomize individuals across chips. For each chip, we freshly prepared a mixture of cell suspensions from four individuals. We measured live cell number via trypan blue staining (ThermoFisher), to ensure equal cell numbers across individuals per mixture. We performed single cell capture and library preparation as previously described using 6 bp Unique Molecular Identifiers [14]. We pooled the 96 samples on each C1 chip and sequenced them on an Illumina HiSeq 2500 using the TruSeq SBS Kit v3-HS (FC-401-3002).

We mapped the reads to human genome GRCh37 (including the ERCC spike-ins) with *Subjunc* [30], deduplicated the UMIs with *UMI*-*tools* [31], and counted molecules per protein-coding gene (Ensembl 75) with *featureCounts* [32]. We then matched single cells back to YRI individuals using *verifyBamID* [33].

We filtered samples on the following criteria, derived as previously described [14]:

- Only one cell observed per well
- Valid identification
- At least 1,011,612 mapped reads
- Less than 49% ERCC reads
- At least 4,730 genes with at least one read
- Linear discriminant analysis predicts one cell

We filtered genes for QTL mapping on the following criteria:

- Number of molecules less than 4^6^ = 4096
- Median log CPM at least 3

We applied principal component analysis (PCA) to the matrix **X** of log counts per million (log CPM), using the pseudocount proposed in *edgeR* [34].

We corrected for gene detection rate by simultaneously regressing out quantiles of gene expression, correcting for sample-specific and gene-specific means, and performing PCA. Let **X** = (**x**_1_,…, **x**_*n*_) be observed *p*-vectors, and let (**z**_1_,…, **z**_*n*_) be latent *k*-vectors where *k* ≪ *p*. Then, PCA corresponds to maximum likelihood estimation in the following latent variable model [35]:

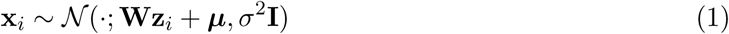

In this parameterization, ***μ*** denotes a per-coordinate mean (in our application, per-gene). However, as previously reported [15], we additionally have to account for the per-sample mean.

Our approach is based on the latent variable model:

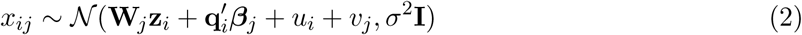

where **u** is an *n*-vector of per-sample means, **v** is a *p*-vector of per-gene means, and **Q** = (**q**_1_,…, **q**_*n*_) is a *k* × *n* matrix of expression quantiles.

We fit the model as follows:

1. Estimate *β_j_* via least squares estimation of the following linear model:

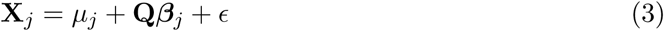
2. Construct the residual matrix
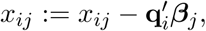
then estimate **u**, **v** via coordinate descent:

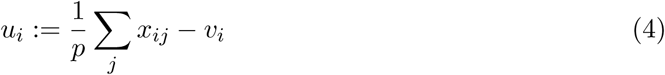

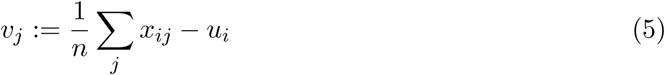
3. Construct the residual matrix *x_ij_*:= *x_ij_* − *u_i_* − *v_j_*, then estimate **W** via maximum likelihood. The MLE **Ŵ** equals the top *k* singular vectors of residual matrix **X** [35].

We estimated the squared correlation between each PC and categorical covariates (batch, C1 chip, individual, well) by recoding each category as a binary indicator, fitting a multiple linear regression of the PC loadings against the binary indicators, and then estimating the coefficient of determination of the model.

### Estimating gene expression mean and variance

We assume the count data are generated by a zero-inflated negative binomial (ZINB) distribution. Let:

- *r_ijk_* be the number of molecules for individual *i*, cell *j*, gene *k*
- *R_ij_* be a size factor for each cell
- *μ_ik_* be proportional to relative abundance
- *ϕ_ik_* be the variance of expression noise
- *π_ik_* be the proportion of excess zeros
- **x***_ij_* be a *q*-vector of confounders per cell
- *β_k_* be a *q*-vector of confounding effects on gene *k*

Then, we assume:

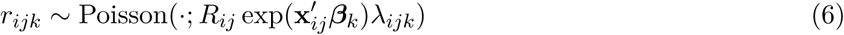

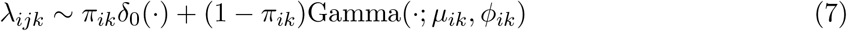

Under this model, the mean and variance of gene expression are:

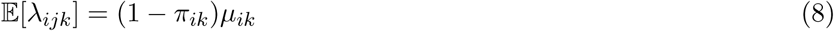

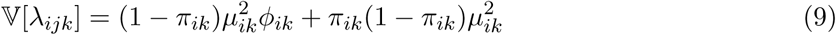

Considering just the non-zero component, marginalizing out λ yields the log likelihood:

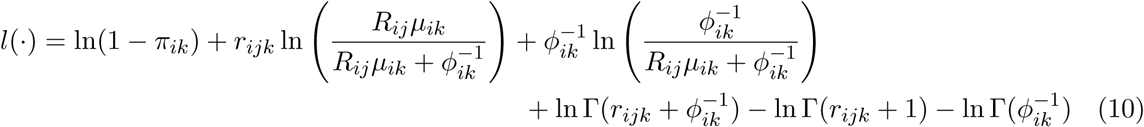

Then, marginalizing over the mixture yields the log likelihood:

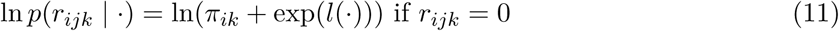

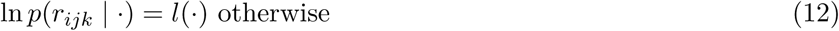

We estimated the model parameters by maximizing the likelihood using batch gradient descent for 4,000 iterations, accelerated by AMSGrad [36].

We defined the size factor of each cell as the total number of molecules detected (before excluding genes in QC). To correct for technical confounders, we included C1 chip as an observed confounder, recoded as binary indicator variables and centered. This approach is sufficient to also correct for batch, because in our experimental design, batch is a linear combination of C1 chip. Intuitively, if there were a batch effect independent of C1 chip, then we could add the batch effect to each chip effect and set the batch effect to 0.

### Quantitative trait locus mapping

We imputed dosages for 120 Yoruba individuals from the HapMap project (Phase 3, hg19) as previously described [37]. We restricted our analysis to 8,472,478 variants with minor allele frequency at least 0.05.

For each single cell expression phenotype tested, we standardized and quantile-normalized the phenotype matrix to a standard normal as previously described [38]. We called QTLs and controlled the gene-level false discovery rate using *QTLtools* [39]. We included principal components (PCs) of the normalized expression matrix as covariates for QTL mapping, and selected the number of PCs for each phenotype by greedily searching for the number of PCs which maximized the number of QTLs discovered on even chromosomes only at FDR 10%. We additionally recalled eQTLs in the matched bulk RNA-seq data [20] using the re-processed dosage matrix.

We performed replication testing by taking each SNP-gene pair from the discovery cohort, and testing that pair in the replication cohort. We defined a hit as replicating if it passed the Benjamini–Hochberg procedure at level 10% (restricted to the set of SNP-gene pairs tested) and had the same effect size direction.

### Power analysis

For individual *i* and gene *k*, we assume the generative model:

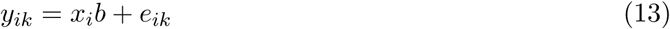

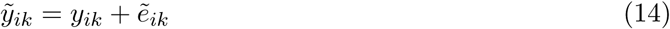

where *ỹ_ik_* is the observed phenotype, *y_ik_* is the true phenotype, *x_i_* is the genotype at the SNP of interest,
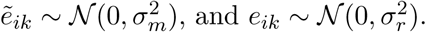

To perform QTL mapping, we fit a working model which ignores measurement error:

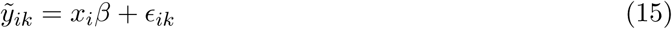

where *ϵ_ik_* ∼ *N*(0, *σ*^2^). From this model, we estimate
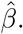
Assuming 𝕍[*x*] = 1, we have
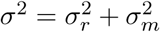
and:

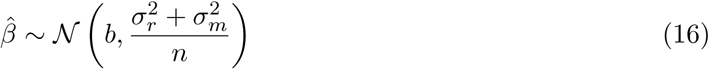

where *n* is the number of individuals. Under the working model, the power function is:

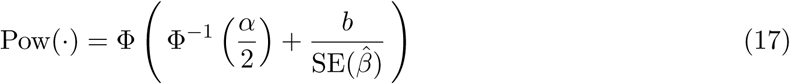

where *α* denotes the significance level, SE(·) denotes standard error, and Φ(·) denotes the standard Gaussian CDF. Under the assumed generative model, the power function equals:

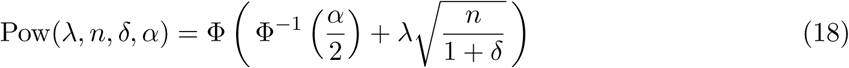

where λ = *b/σ_r_*, and
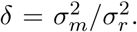
Parameterized in terms of *δ*, the power function implies useful reference points; for example, *δ* = 1 is equivalent to cutting the sample size in half.

To determine the effect size *b*, we estimate the distribution of true effect sizes *b* given observed effect sizes
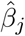
and associated standard errors *ŝ_j_*. We assume the hierarchical model:

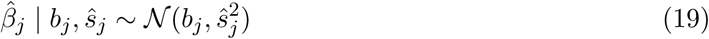

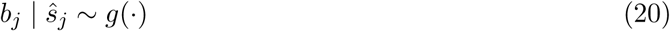

where *g* is a unimodal mixture of Gaussians. We estimate *g* using adaptive shrinkage (*ash*) [21]. We took *b* to be the 99^th^ percentile of the fitted distribution.

Although we assumed a single measurement error variance
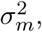
we actually have measurement errors for each individual and gene
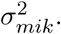
To estimate
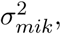
we used non-parametric bootstrapping. For each individual and gene, we resampled the counts (matched with the library size and technical confounders) with replacement, and refit the ZINB model. To reduce computational burden, we restricted our analysis to 200 randomly chosen genes, warm-started the optimization from the optimal parameters for the original data, and ran gradient descent for 1,000 iterations.

To estimate the typical noise ratio *δ*, we estimate a measurement error variance per gene
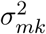
and a residual variance per gene
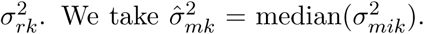
To estimate
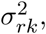
we solve a deconvolution problem [40]:

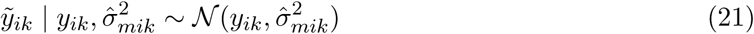

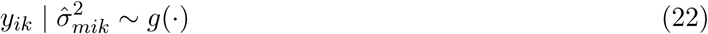

where *g* is a unimodal mixture of uniforms, estimated using *ash*. To fit the model, we centered the *ỹ_ik_* for each gene *k*, concatenated them across genes, and assumed a common prior.

Then, the required estimates are:

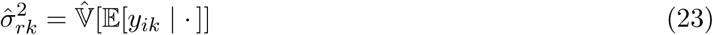

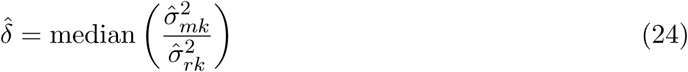

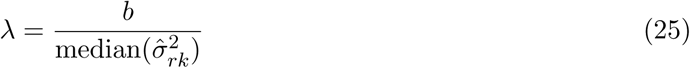

where
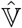
denotes sample variance.

### Code and data availability

All code used to perform the computational analyses is available at https://github.com/jdblischak/singlecell-qtl. The results of running the analysis code are available at **https://jdblischak.github.io/singlecell-qtl**.

The RNA-seq data, sample metadata, and filtered gene expression count matrix have been deposited under accession number GSE118723. The estimated parameter matrices and QTL summary statistics are available at https://eqtl.uchicago.edu

## Acknowledgements

We thank Joyce Hsiao for helpful discussions. Computing resources were provided by the University of Chicago Research Computing Center. This study was supported by NIH grant RO1GM122930.

## Supporting information

**Figure S1:**
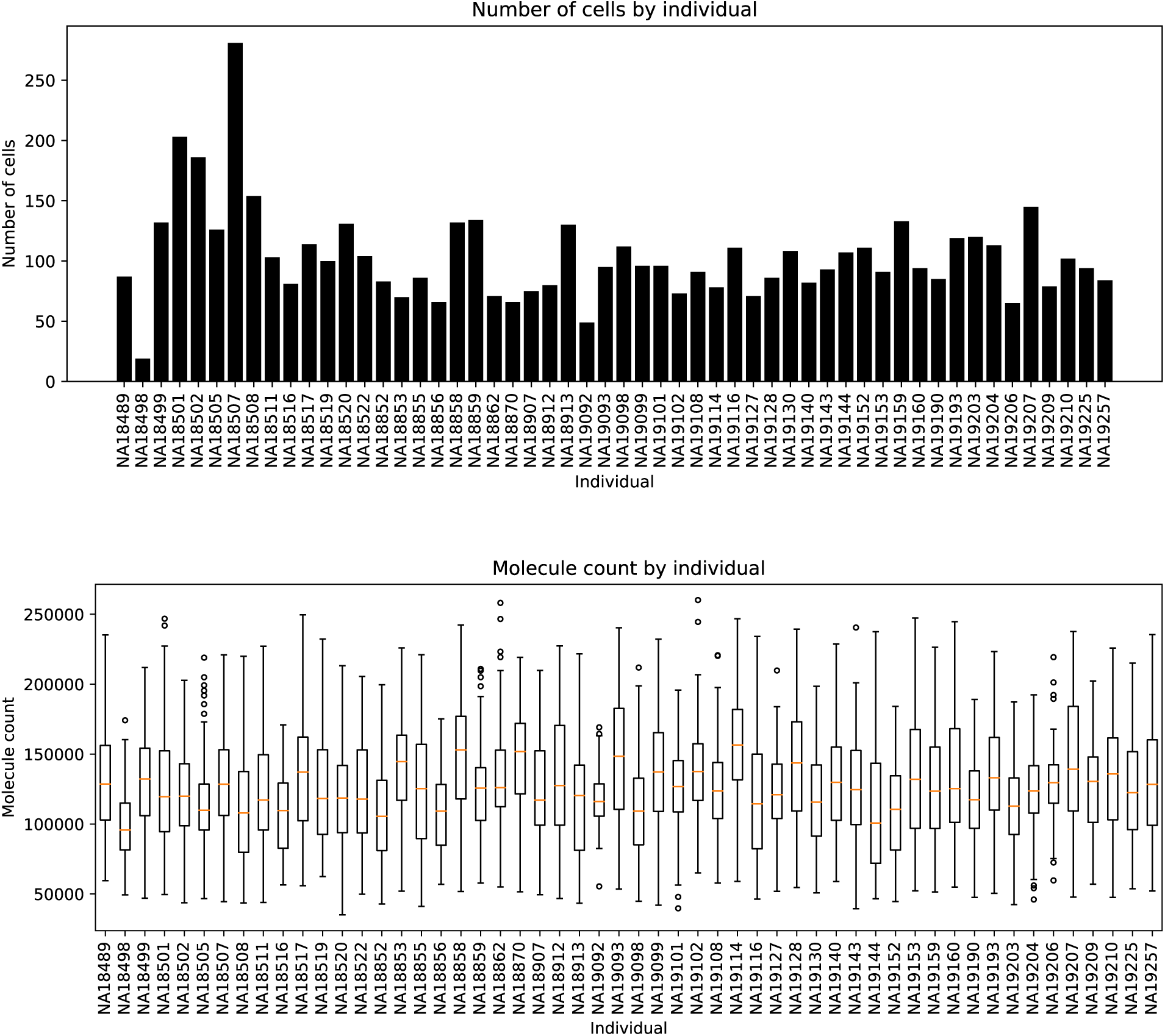
Descriptive statistics of the experiment. Number of cells per individual, and number of molecules per cell after applying quality control filters.

**Figure S2:**
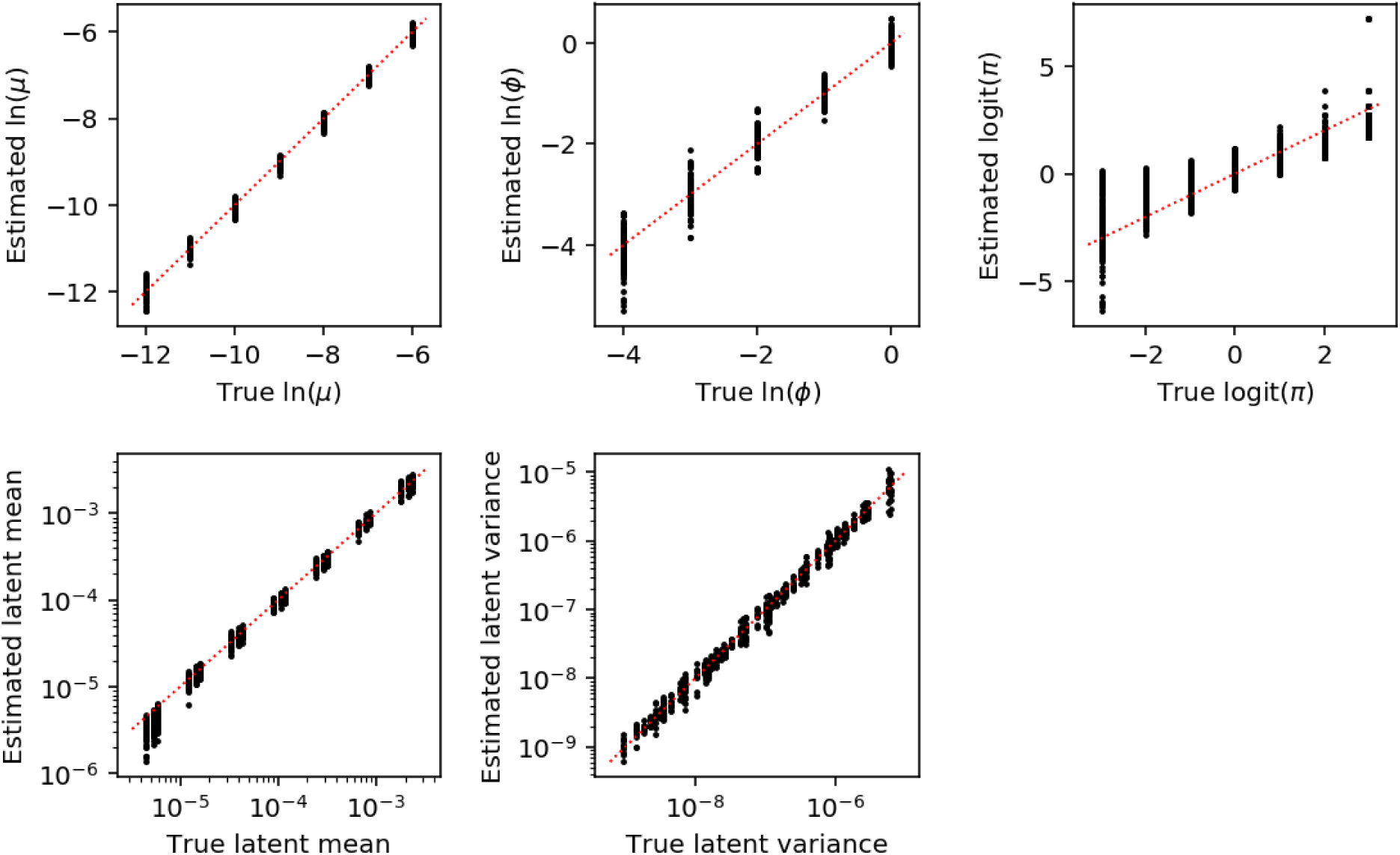
Estimated ZINB parameters and latent mean and variance in idealized simulation. Estimates of ln(*μ*) and latent mean are displayed for logit(*π*) < 0. Estimates of ln(*ϕ*) and latent variance are displayed for ln(*μ*) > −10, logit(*π*) < 0. Estimates of logit(*π*) are displayed over the entire range of parameter values. In each trial, simulated molecule counts for 95 cells are drawn from the model assuming 114,026 molecules per cell, matching the median number of cells, and molecules per cell in the observed data.

**Figure S3:**
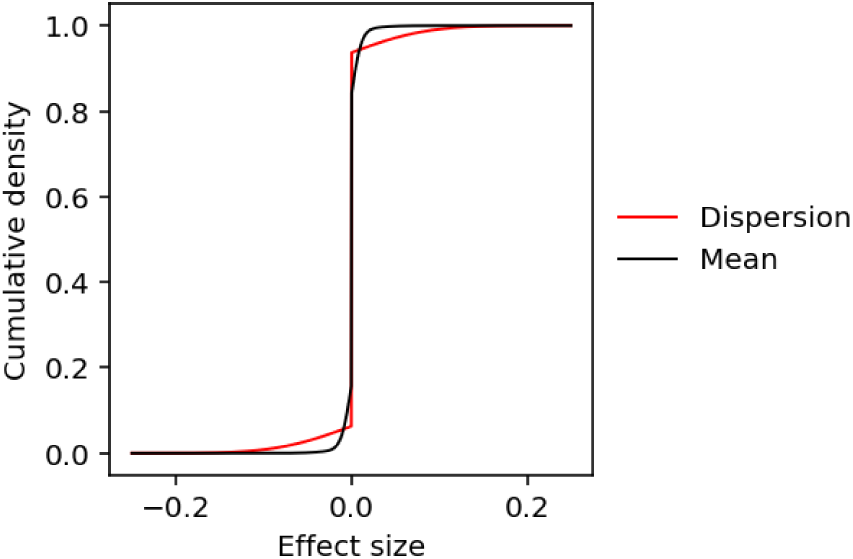
Estimated distribution of QTL effect sizes. We fit a unimodal mixture of Gaussians to the distribution of observed eQTL (dQTL) effect sizes (in terms of log fold change) using Empirical Bayes.

**Figure S4:**
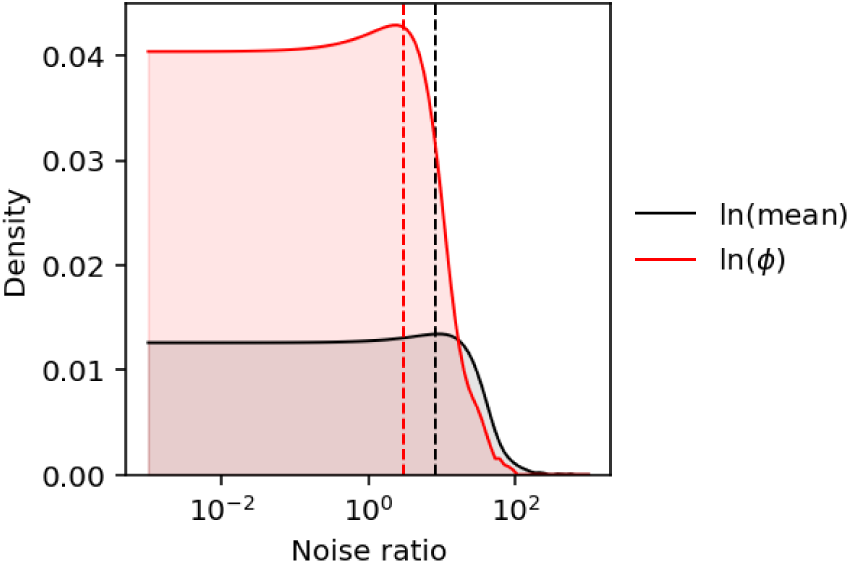
Distribution of estimated noise ratios. Noise ratios (ratio of measurement error variance to phenotypic variance) are estimated for 200 randomly chosen genes using a two-step empirical Bayes procedure.

